# Fifty shades of white fish: DNA barcoding reveals widespread misidentification in sushi restaurants from Southern Brazil

**DOI:** 10.1101/2025.05.02.651977

**Authors:** Fernanda Almerón-Souza, Leonardo Tresoldi Gonçalves, Nelson Jurandi Rosa Fagundes

## Abstract

The ambiguous commercial naming of fish can mislead consumers and obscure species identity. “White fish” is a widely used yet unregulated commercial name in sushi restaurants across Brazil, creating opportunities for species misidentification. This study used DNA barcoding to assess the accuracy of species labeling for “white fish” sushi in Porto Alegre, Southern Brazil, a major urban center with a high density of sushi restaurants. A total of 53 “white fish” sushi samples were collected from 50 restaurants. Molecular identification successfully assigned 41 samples to nine distinct fish species, while seven samples were identified at the genus level. In eight cases (20.5%), the species name provided by restaurant staff did not match the molecular identification, meaning that one in five restaurants supplied incorrect information. We found that lower sushi prices were significantly associated with a higher likelihood of misidentification (*P*=0.035, OR = 0.53, 95% CI: [0.17, 1.03]). Among the eight cases of species substitution, three included the substitution of “*linguado*” (flounder; *Paralichthys* spp. or *Syacium* spp.) for “*panga*” (striped catfish; *Pangasianodon hypophthalmus)*, and another three revealed the substitution of “*prego*” (escolar; *Lepidocybium flavobrunneum* or *Ruvettus pretiosus*) for “*meca*” (swordfish; *Xiphias gladius*). The remaining two cases revealed the swapping between “*tilápia*” (tilapia; *Oreochromis* spp.) and “*prego*” (escolar). These findings highlight the need for stricter seafood labeling regulations and enforcement to improve consumer transparency and sustainability.

**Highlights:**

- “White fish” corresponded to at least ten different fish species;
- One in five restaurants provided incorrect information regarding the fish species identification;
- High-value species were replaced with less economically valuable ones;
- Lower sushi prices were associated with a higher likelihood of misidentification;
- Brazilian legislation should regulate the commercial use of the term “white fish”.

## 1. Introduction

Fish meat is among the most relevant food resources worldwide and one of the most traded commodities. In recent decades, global aquaculture and fisheries production has grown by an average of 3.2% annually, with demand for fish expected to continue rising (FAO, 2018). However, large-scale exploratory fishing without adequate conservation measures has led to major declines in fish populations (Barletta et al., 2010; FAO, 2018). This has resulted in fluctuations in fish quality, resource availability, and market stability, leading to price volatility (Hellberg & Morrissey, 2011). The combination of increasing global fish consumption, the rise in processed food production, and price disparities among different fish species has created an environment conducive to economic fraud in the food industry, particularly in the form of species substitution (Fox et al., 2018; Hellberg & Morrissey, 2011).

Sushi, a traditional Japanese dish made with vinegared rice and typically topped or filled with raw seafood, has become a widely favored food around the world. Factors such as global trade, health-conscious eating trends, and the simplicity of its preparation have contributed to its international popularity (Edwards, 2012). In Japan, a wide variety of fish species are traditionally used in sushi preparation (Quaas & Requate, 2013). However, a diverse set of species is used in different countries depending on their availability. In Brazil, for example, sushi restaurants often limit their offerings to tuna, salmon, and “white fish”, a generic descriptive commercial term. While Brazilian food legislation strictly links “*atum*” (tuna) and “*salmão*” (salmon) to eight and six specific species, respectively, “peixe branco” (“white fish”) lacks an official definition (Instrução Normativa MAPA N° 29, 2015; Instrução Normativa MAPA N° 53, 2020; Portaria MAPA N° 570, 2023), allowing any fish with relatively white flesh to be marketed under this label. This regulatory gap not only obscures the identity of the fish being sold but also makes it difficult for consumers to know what they are actually eating.

Since sushi involves cutting the fish into pieces, it becomes impossible for consumers to identify the species, which typically relies on the external morphology of the whole fish (Vandamme et al., 2016). This scenario allows for both accidental and intentional species substitution, resulting in consumer deception and unawareness, potential health risks from consuming certain fish species, and environmental issues related to illegal or unregulated fishing (Lowenstein et al., 2010; Quaas & Requate, 2013). For example, substituted species may contain higher levels of toxins such as mercury or allergens, posing direct health risks to consumers (Marko et al., 2014; Triantafyllidis et al., 2010). Moreover, the use of illegally caught or overfished species undermines conservation efforts and sustainable fishing practices, further intensifying environmental degradation (Almerón-Souza et al., 2018; Yan et al., 2021).

DNA barcoding is a molecular biology technique that uses short, standardized DNA sequences to identify species by comparing them to reference databases (Hebert et al., 2003). In animals, the most widely used DNA barcode is a 650-base pair (bp) fragment of the mitochondrial gene cytochrome *c* oxidase subunit I (cox1 or COI). This method has become a cornerstone in species identification studies, prompting many countries to adopt legislation integrating it into seafood traceability systems (Clark, 2015; Handy et al., 2011). Although Brazil lacks specific regulations for its use in seafood trade, DNA barcoding has gained traction in detecting species substitution, particularly during periods of heightened demand for specific fish, such as cod during the Easter and Christmas seasons (Calegari et al., 2020; Carvalho et al., 2015). However, no study has explored which species represents “white fish” in sushi restaurants in Brazil, nor if restaurants are able to inform this correctly.

In this study, we utilized DNA barcoding to identify the species being sold as “white fish” across the city of Porto Alegre, a major urban center in Southern Brazil with a high density of sushi restaurants. We then compared the DNA-based identifications with the commercial names provided at the time of purchase to assess the accuracy of species labeling. Additionally, we tested whether sushi price and neighborhood socioeconomic status influenced the risk of species misidentification. Our findings are discussed under the light of the regulatory gaps in Brazil’s seafood labeling practices, raising awareness about the broader implications of species substitution.

## 2. Material and methods

### 2.1. Sampling

From February to June 2019, we used food delivery apps to search for sushi restaurants located in Porto Alegre, Rio Grande do Sul, a major urban center in Southern Brazil. Whenever we found items being sold under the terms “*peixe branco*” (“white fish”) or “*sushi branco*” (“white sushi”), the item was purchased (e.g. a set of *sashimi* or *nigiri* pieces), and a phone call was made to each restaurant to inquire about the fish species used for the “white fish” sushi.

Overall, we ordered 53 samples labeled as “white fish” from 50 sushi restaurants. The common names of the fish provided at the time of purchase were recorded, along with the price per sushi piece, and other data regarding the restaurant. Information about fish suppliers was requested upon a second contact from those restaurants whose samples were later found to have been incorrectly labeled, but without revealing the results from DNA identification. We also contacted nearly all remaining sushi restaurants in Porto Alegre to ask whether they sold “white fish” sushi, to ensure that the sample collection was as broad as possible and covered different neighborhoods and price ranges.

The samples consisted of “white fish” sushi pieces in the forms of *nigiri, sashimi*, and *hosomaki*. For *nigiri* and *hosomaki*, the rice was removed before freezing the fish at -4ºC. A tissue fragment (approximately 25 μg) was taken from the inner portion of each sample to minimize the risk of contamination from handling or knife-sharing at the restaurant, and was then stored in 95% ethanol.

### 2.3. DNA extraction, amplification, and sequencing

The samples were thawed and then homogenized. DNA extraction was performed using the CTAB method (Doyle & Doyle, 1987). Quality control of the extraction was carried out using a NanoDrop 8000 spectrophotometer, which measured both the concentration and purity of the DNA samples. Polymerase Chain Reaction (PCR) was employed for DNA amplification, using primer pairs FishF2 (5’ TCGACTAATCATAAAGATATCGGCAC 3’) and FishR2 (5’ ACTTCAGGGTGACCGAAGAATCAGAA 3’) (Ward et al., 2005), designed to amplify a fragment of approximately 650 bp of the mitochondrial COI gene. The PCR reaction was performed in a final volume of 20 μL, containing 0.4 μM of each dNTP, 1.5 mM of MgCl_2_, 0.25 μM of each primer, 1U of Taq polymerase, and 40 ng of DNA. The amplification process began with an initial denaturation at 94°C for 5 minutes, followed by 10 cycles of 94°C for 1 minute, 55°C (with a 0.5°C decrease per cycle) for 1 minute, and 72°C for 1 minute and 30 seconds. This was followed by 30 additional cycles of 94°C for 1 minute, 50°C for 1 minute, and 72°C for 1 minute and 30 seconds, with a final extension step at 72°C for 5 minutes. Quality of PCR products was assessed by 1% agarose gel electrophoresis stained with GelRed. We purified PCR products using exonuclease I and shrimp alkaline phosphatase. DNA sequencing was conducted by ACTGene Molecular Analyses (Porto Alegre, Brazil) in both directions using the Sanger method.

### 2.4. Sequence analysis and sample identification

We used Geneious (http://www.geneious.com/) to visually inspect chromatograms and generate consensus sequences. For species identification, we employed two complementary methods. First, we used the Barcode of Life Data System (BOLD; Ratnasingham et al., 2024) identification workbench available at https://id.boldsystems.org/, querying the sequences against BOLD’s Animal Library, that includes over 7.5 million non-redundant COI sequences of at least 500 bp within the barcode region. This system returns a species-level identification including a confidence value and a list of BOLD records supporting the identification. Second, we used CO1 Classifier v5.1.0 (Porter & Hajibabaei, 2018), a training set that can be used with the Ribosomal Database Project (RDP) classifier (Wang et al., 2007) to taxonomically assign COI sequences. This method circumvents issues that arise when using NCBI’s BLASTn for direct comparisons with GenBank data, including the high false positive rate and the lack of a measurement of statistical confidence in the taxonomic assignments (Porter & Hajibabaei, 2018).

### 2.5. Evaluation of the commercial names informed by restaurants

To assess whether the fish name provided at the time of purchase corresponded to the species identified using DNA barcoding, we utilized the list of commercially relevant fish from the Brazilian Ministry of Agriculture’s Normative Instruction 29/2015, which was in vigor at the time of sample collection (Instrução Normativa MAPA N° 29, 2015). This list associates common fish names with the species that can be legally sold under each common name. Notably, the term “*peixe branco*” (“white fish”), widely used in sushi restaurants, is not recognized as an official common name in the Brazilian regulatory list. Therefore, we focused on the common names verbally provided by the establishments and considered that a correct identification occurred whenever the scientific name identified in the molecular analysis matched the commercial name reported by the restaurant, in accordance with the legally recognized commercial names specified in the abovementioned normative instruction. For example, if the restaurant informed that the items sold corresponded to “*linguado*” (flounder), any species belonging to *Paralichtys* spp. or *Syacium* spp. would correspond to a correct identification. Additionally, we cross-referenced species identifications using FishBase (http://fishbase.se) to validate scientific nomenclature and assess the alignment between the common names listed in the Normative Instruction and those used in scientific contexts.

To evaluate putative predictors of mislabeling, we built a logistic regression model to evaluate the relationship between misidentification (binary dependent variable) and the independent variables, which included sushi price per piece (continuous) and the socioeconomic profile of the neighborhood (continuous, represented by the average household income, in minimum wage units per month; Porto Alegre, 2019). The model was fitted using the glm() function in R v.4.3.2 (R Core Team, 2023) with a binomial family and logit link function. The significance of the model and its predictors was assessed using analysis of deviance with a Chi-square test. Model fit was evaluated using McFadden’s pseudo-*R*^*2*^, calculated using the pscl R package (Jackman, 2005), with values around 0.2–0.4 considered an excellent fit (McFadden, 1979). Three samples were excluded from the logistic regression: sample 43, as it was obtained from an all-you-can-eat buffet and no unit price of sushi was available; and samples 3 and 27, which were identified as bacterial sequences (see Section 3.2).

## 3. Results and discussion

### 3.1. Data collection

Out of the total 124 sushi restaurants surveyed, 63 reported offering “white fish” sushi on their menu. No samples were collected from 13 of these establishments because in three, “white fish” sushi was temporarily unavailable, while in the remaining 10, staff indicated that tilapia was used to prepare the sushi. Given that tilapia in Brazil is a farmed and relatively inexpensive fish, and multiple samples had already been purchased under the commercial name of tilapia, priority was given to samples labeled as other species.

Only three establishments listed a specific market name of fish on their menus instead of using the generic term “white fish”. In other locations, this information could only be obtained by asking the restaurant staff. Although we were able to gather information from all sampled establishments about the species marketed as “white fish”, this was sometimes challenging, as staff were not always aware of the fish species used and had to consult with kitchen personnel. Some restaurants reported that they did not have “white fish” available, indicating a preference for specific white-fleshed fish species. Others noted that they served different species as “white fish” depending on the day or week, suggesting that there is considerable variability in the types of fish offered under this designation.

Interestingly, despite the general understanding that “white fish” refers to white-fleshed species, some fish sold under this term had pink flesh (Fig. 1), often sold as swordfish (*Xiphias gladius*, “*meca*”). One restaurant, though not offering “white fish” sushi, provided swordfish *nigiri*, noting that swordfish has pink flesh rather than white. This situation shows how the lack of specific regulations for selling “white fish” in Brazil leads to a broad interpretation of the term, allowing sushi restaurants to market various pale-fleshed species (especially in comparison to common species such as salmon and tuna) as “white fish”.

**Fig. 1.**
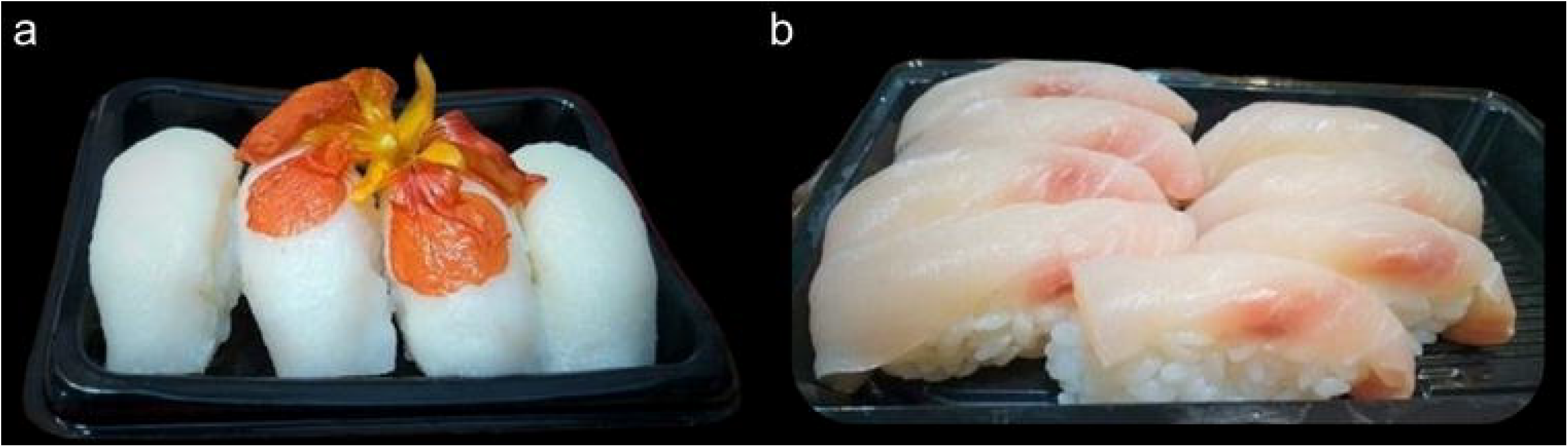
Samples collected as “white fish” *nigiri*. (a) Sample sold as “prego” (escolar, *Lepidocybium flavobrunneum*), with a very white flesh. (b) Sample sold as “meca” (swordfish, *Xiphias gladius*), exhibiting a pinkish hue. Samples are shown as sent, including a decorative *Delonix regia* flower in photo “a”.

### 3.2. Molecular identification of samples

A total of 41 samples were successfully amplified and sequenced. Of these, 32 samples were identified to the species level, representing nine distinct fish species: *Xiphias gladius* (swordfish, *n* = 11), *Lepidocybium flavobrunneum* (escolar, *n* = 7), *Paralichthys orbignyanus* (Brazilian flounder, *n* = 5), *Pangasianodon hypophthalmus* (striped catfish, *n* = 3), *Paralichthys patagonicus* (Patagonian flounder, *n* = 2), *Coryphaena hippurus* (common dolphinfish, *n* = 1), *Pagrus pagrus* (Red porgy, *n* = 1), *Pseudopercis semifasciata* (Argentinian sandperch, *n* = 1), and *Seriola rivoliana* (longfin yellowtail, *n* = 1). All obtained identifications had a minimum BOLD confidence score of 91 and a CO1 Classifier bootstrap value of at least 0.92 (Table 1; Table S1; Table S2). Importantly, the species identifications provided by the BOLD database and the CO1 Classifier were fully concordant (Table S1; Table S2).

**Table 1.**
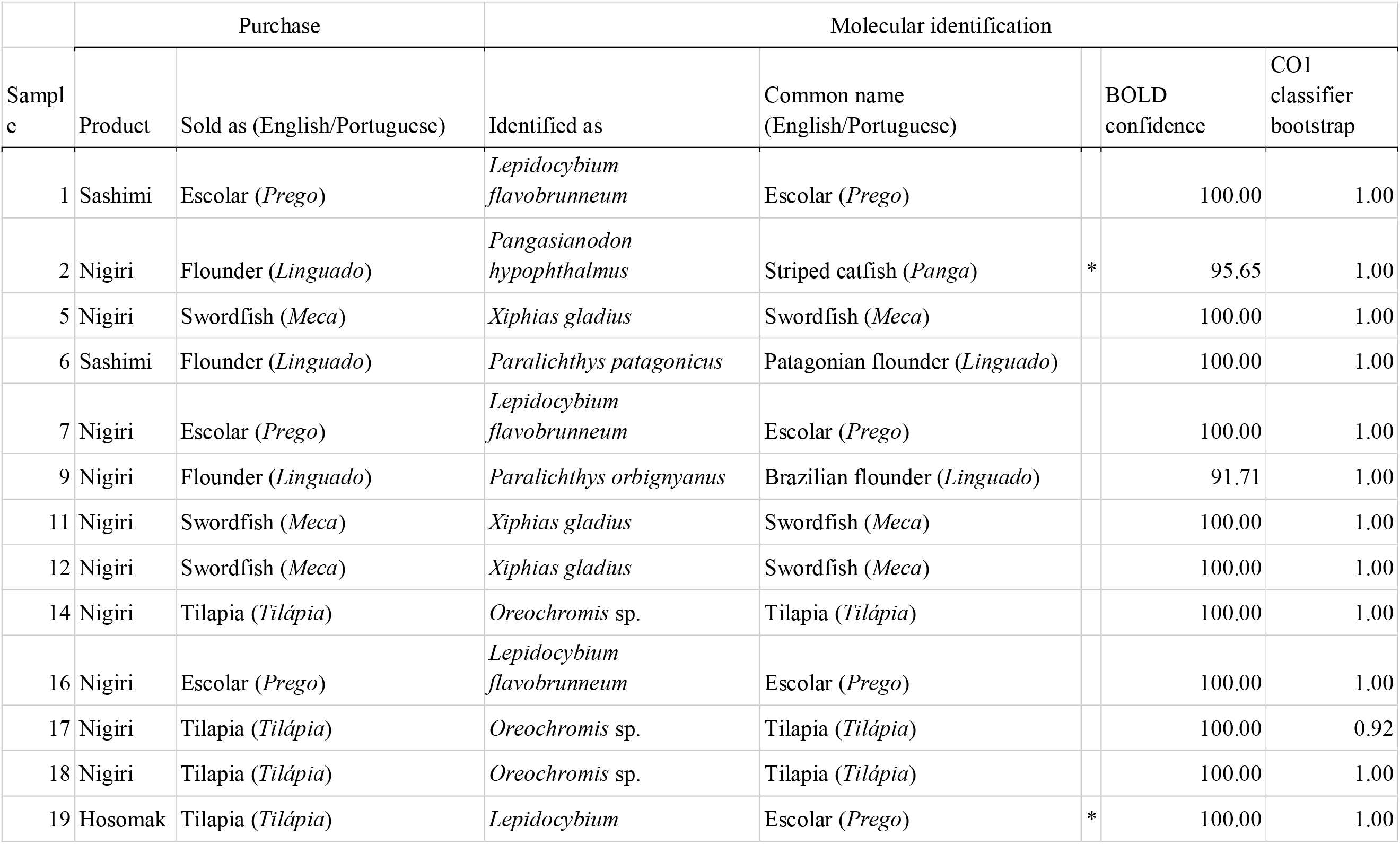
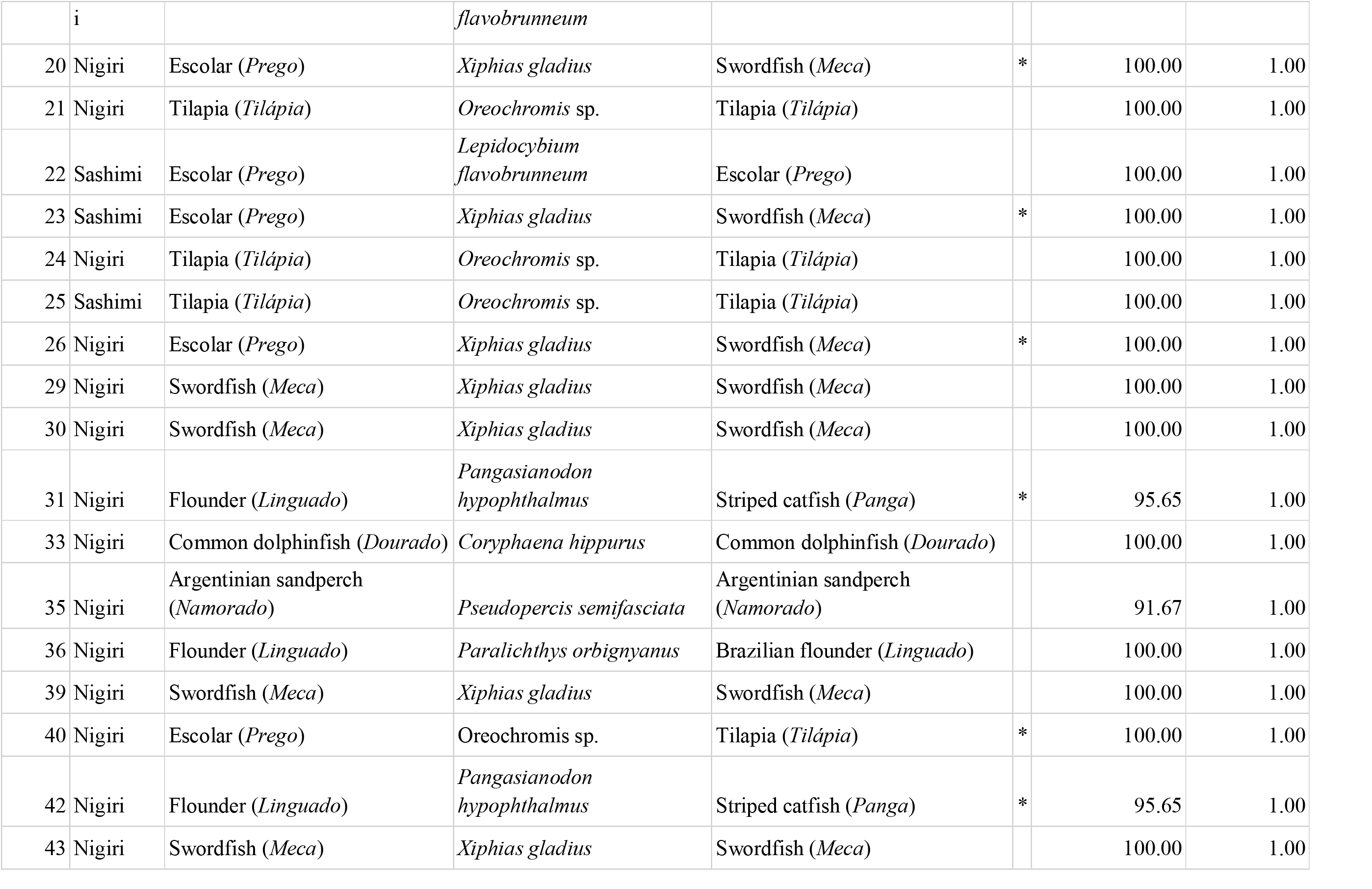
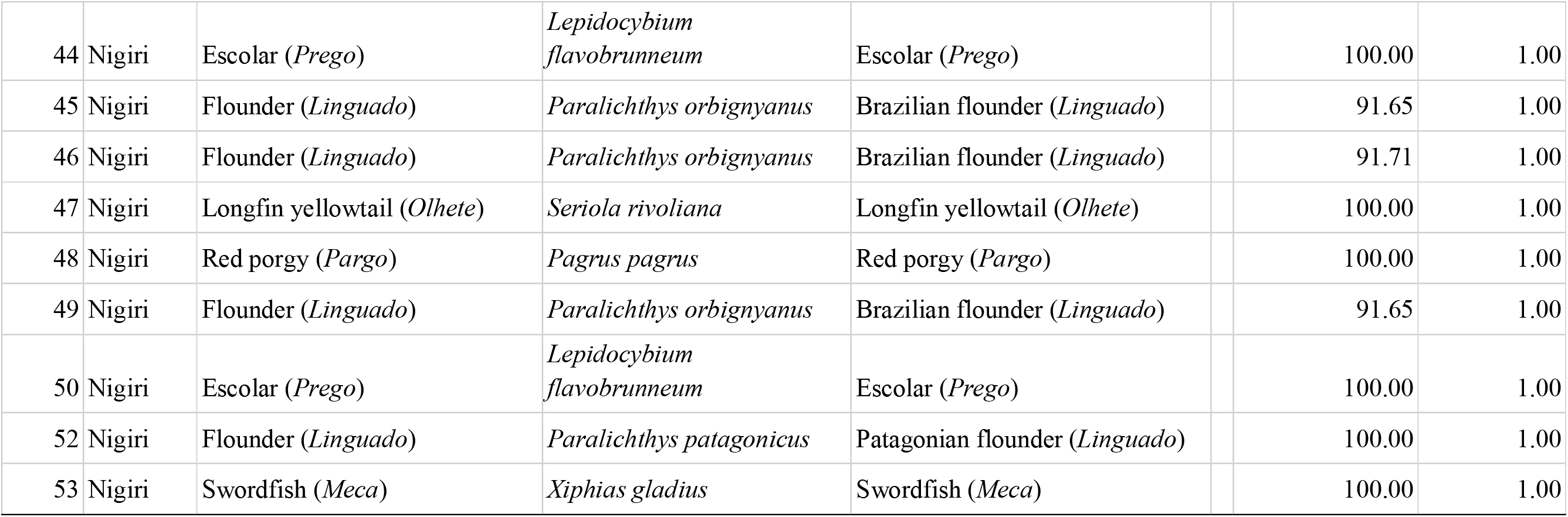
Summary of the 42 samples analyzed in this study. Lines marked with an asterisk indicate mismatch between species informed during purchase and molecular identification. Support values for samples identified as *Oreochromis* sp. refer to genus-level placement.

Seven samples were identified at the genus level as *Oreochromis* sp. (tilapia). The inability to resolve these samples to the species level is likely attributable to the long history of hybridization practices in tilapia farming (Mojekwu et al., 2021; Nascimento et al., 2023; Pollack et al., 2018). Nonetheless, both BOLD and CO1 classifier results confirmed the genus-level assignment as *Oreochromis* with high support for all these samples (Table 1).

Two samples (3 and 27; sold as “*escolar*”, *L. flavobrunneum*) yielded high-quality sequences that matched the COI-like gene of a bacterium from the genus *Pseudomonas*, specifically *P. fragi*. This result persisted even after repeating the DNA extraction, PCR amplification, and sequencing processes using the original samples. As expected, no matches for these sequences were found in the BOLD database, as it is restricted to eukaryotic taxa. However, the CO1 classifier assigned these sequences to *P. fragi* with high bootstrap support (BS = 1; Table S2). *Pseudomonas fragi* is known to be associated with the spoilage of seafood, including refrigerated fish (Darwish et al., 2023; Gram et al., 2002; Luqman et al., 2024). Potential origins for such spoilage may include improper handling during preparation, extended storage prior to processing, or opportunistic bacterial growth on the fish (Ardura et al., 2013). In general, the presence of *Pseudomonas* spp. in food is not considered as a serious risk to public health (Raposo et al., 2017). However, the presence of antibiotic-resistant strains and of the human pathogenic species *P. aeruginosa* in other studies begs for good hygienic measures in the handling of raw fish food products (Cordeiro et al., 2020; Darwish et al., 2023).

### 3.3. Mislabeling and its economic and regulatory implications

The molecular analysis revealed that of the 39 samples analyzed, 31 (79.5%) matched the vernacular names provided by restaurants, while eight samples (20.5%) were mislabeled(Table 1, Fig. 2). This means that in Porto Alegre one in five sushi restaurants provided incorrect species names when asked about the identity of the species sold as “white fish”. Interestingly, the restaurants reported buying fish from common suppliers that operate in other regions of Brazil (data not shown), suggesting the issue extends beyond local markets.

**Fig. 2.**
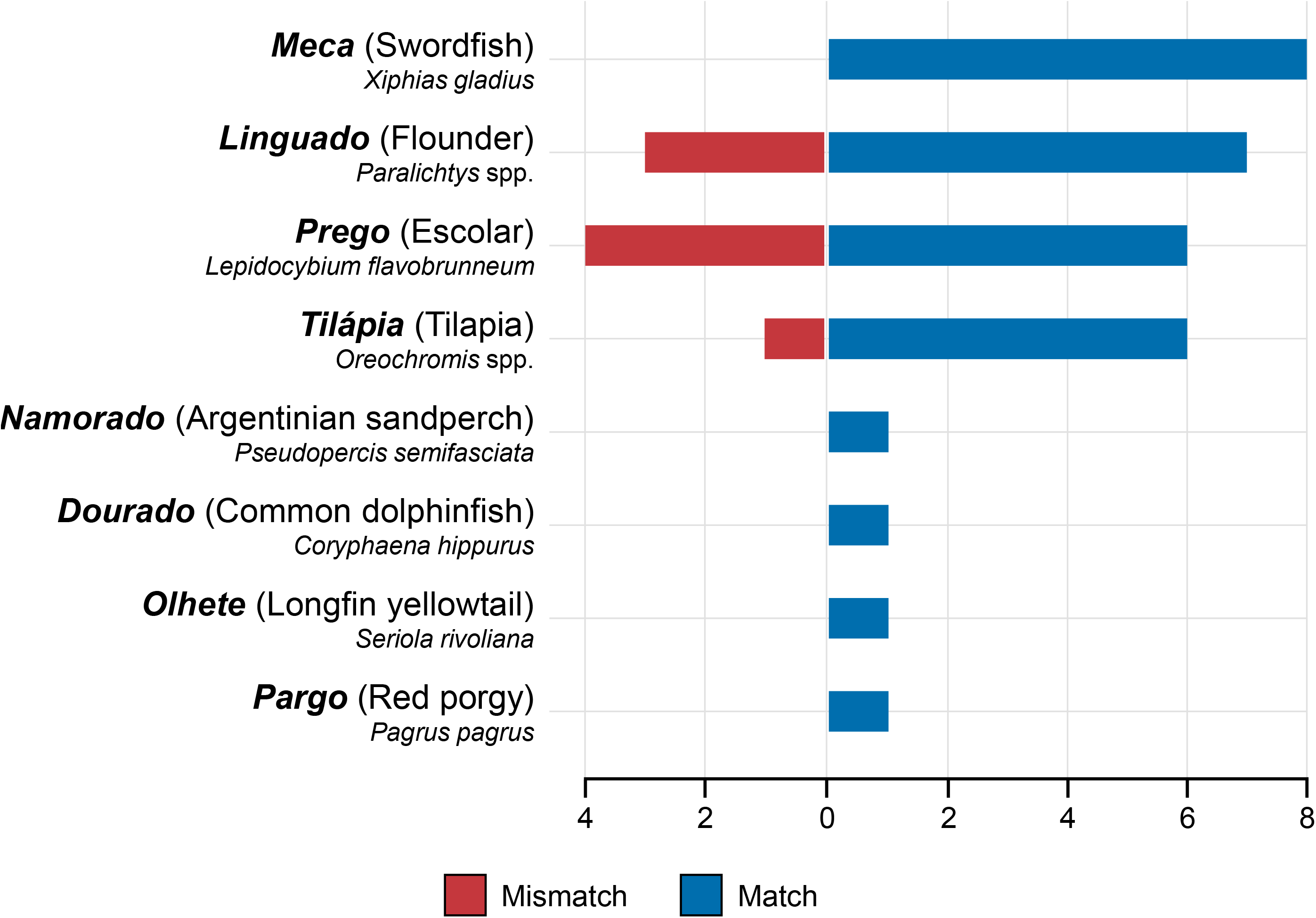
Relationship between molecular identification results and the commercial names provided by sushi restaurants. Matches (blue, right side) and mismatches (red, left side) are shown for each vernacular name. Importantly, Brazilian regulations also permit the use of “linguado” for *Syacium* spp., “prego” for *Ruvettus pretiosus*, “namorado” for *Pseudopercis numida*, “dourado” for *Salminus* spp., “olhete” for *Seriola* spp., and “pargo” for *Lutjanus* spp. (Instrução Normativa MAPA N° 29, 2015).

For the eight mislabeled items, the most common cases of species substitution affected three samples (37.5% of the mislabeled items), and included the substitution of items sold as “*linguado*” (flounder; *Paralichthys* spp. or *Syacium* spp.) that corresponded to “*panga*” (striped catfish; *Pangasionodon hypophtalmus*). Another three items revealed the substitution of “*prego*” (escolar; *L. flavobrunneum* or *R. pretiosus*) for “*meca*” (swordfish; *X. gladius*). The remaining two misidentified samples (25% of the mislabeled items) involved the same pair of species, “*tilápia*” (tilapia; *Oreochromis* spp. and hybrids with *Sarothedon galilaeus*) and “*prego*” (escolar), but while in one case the item was sold as tilapia and corresponded do escolar, the opposite occurred in the other case. In all cases, mislabeling occurred between species with marked morphological differences, making accidental mislabeling unlikely (Fig. 3), and in five cases (62.5% of the mislabeled items) it involved marine (flounder and escolar) and freshwater (striped catfish and tilapia) species, increasing the possibility of a deliberate substitution.

**Fig. 3.**
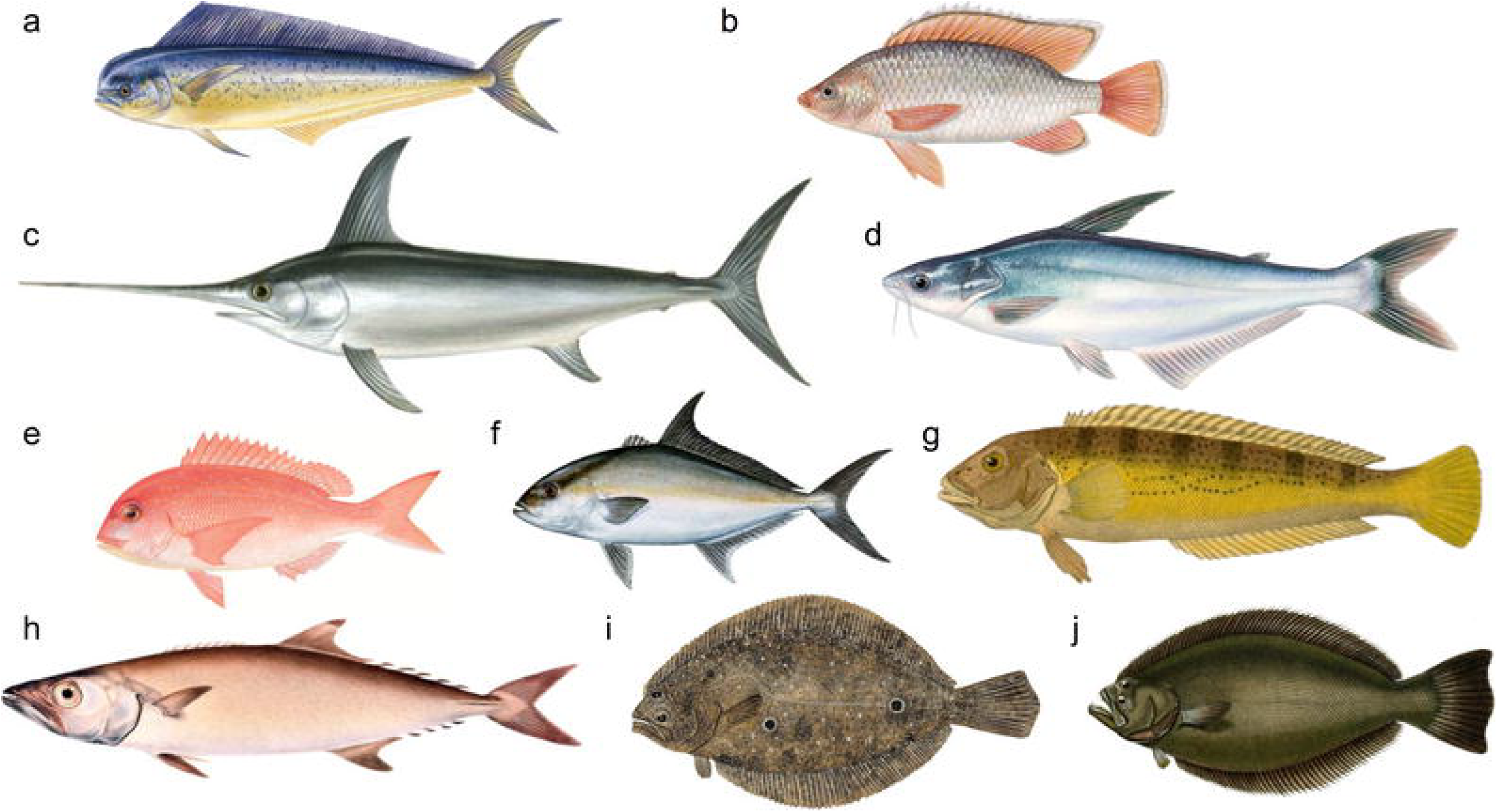
Fish species identified in “white fish” sushi using DNA barcoding in this study. The species exhibit significant morphological diversity, making it highly unlikely that substitutions occurred due to misidentification. (a) Common dolphinfish (*Coryphaena hippurus*); (b) Tilapia (*Oreochromis niloticus*); (c) Swordfish (*Xiphias gladius*); (d) Striped catfish (*Pangasianodon hypophthalmus*); (e) Red porgy (*Pagrus pagrus*); (f) Longfin yellowtail (*Seriola rivoliana*); (g) Argentinian sandperch (*Pseudopercis semifasciata*); (h) Escolar (*Lepidocybium flavobrunneum*); (i) Patagonian flounder (*Paralichthys patagonicus*); (j) Brazilian flounder (*Paralichthys orbignyanus*). Fish are depicted out of scale. Pictures sourced under a Creative Commons license. Credits: a, b, c, e, Scandinavian Fishing Year Book; d, WWF; f, Diane Rome Peebles, g, Cuvier et al. (1828); h, Smith et al. (1838); i, Kim In Young; j, Paul Louis Oudart.

A logistic regression model indicated that lower sushi prices were significantly associated with higher odds of misidentification (*P* = 0.035, OR = 0.53, 95% CI: [0.17, 1.03]), such that each unit decrease in price was associated with an 89% increase in the odds of misidentification. In contrast, neighborhood income had no significant effect (*P* = 0.377, OR = 0.88, 95% CI: [0.64, 1.16]) (Table S3). The model achieved a McFadden’s pseudo-R^2^ = 0.13, indicating modest improvement over the null model. The substitution of higher-value species with lower-value ones is a well-documented market practice driven by economic incentives, especially when the items are sold in the absence of morphological indicators that are species specific (Kappel & Schröder, 2016). The most significant price discrepancy observed was in the substitution of “*linguado*” (flounder) with “*panga*” (striped catfish), as, at the time of sampling, the kilogram price of flounder was three times more expensive than of striped catfish. Similar cases of *P. hypophthalmus* mislabeling have been reported worldwide, including its sale as cod in Canada, Egypt, and Europe (Hanner et al., 2011; Galal-Khallaf et al., 2014, Pardo et al., 2018), as grouper and other percids in Europe (Di Pinto et al., 2015; Pardo et al., 2018), and as flounder in Southern Brazil (Carvalho et al. 2015). While fish prices appear to be a key factor affecting species substitution, additional data will be needed to test for other predictive variables, such as seasonality.

The mislabeling rate detected in this study (25.6%) aligns with previous research in Brazil using DNA barcoding. Two studies conducted in Florianópolis, a major tourist hub in Southern Brazil, found mislabeling values between 24–26% (Carvalho et al., 2015; Staffen et al., 2017). A national-wide survey found a slightly lower mislabeling value of 17.3%, but a higher (though non-significant) incidence in Southern Brazil (Carvalho et al., 2017). Internationally, a systematic review of 51 fish authentication studies found an average mislabeling rate of 30%, though rates varied widely from 0% to 84% (Pardo et al., 2018). However, differences in sampling methodologies, such as whether studies targeted specific stages of the supply chain (restaurants, fish markets, or suppliers) or included products already suspected of fraud, make challenging a straightforward comparison of different studies.

Even when fraud is not apparent, restaurant nomenclature can obscure species identity. Brazilian fish naming conventions often assign a single commercial name to multiple species, genera, or even families (Instrução Normativa MAPA N° 29, 2015), limiting consumer transparency (Cawthorn et al., 2018; Xiong et al., 2016). A notable example is “*cação*”, commonly used in Brazil to refer to any shark or ray species, though even bony fish species end up marketed under this name (Almerón-Souza et al., 2018). Indeed, most names provided by restaurants were “umbrella” popular names encompassing multiple species within or across genera. For instance, “*tilápia*”, “*namorado*”, and “*pargo*” correspond to multiple species within *Oreochromis, Pseudopercis*, and *Lutjanus*, respectively, while “*prego*”, “*linguado*”, and “*dourado”*“span different genera. To improve transparency, each traded species should be assigned a unique commercial name, and scientific names should be mandated at some point of the processing chain—an approach already required for species within the families Salmonidae (salmonids) and Gadidae (true cods) (Instrução Normativa MAPA N° 29, 2015). Such measures are critical for ensuring consumers receive accurate information about the fish they purchase. Subsequent regulations for fish labeling practices maintain the use of umbrella terms in Brazil (Instrução Normativa MAPA N° 53, 2020; Portaria MAPA N° 570, 2023).

### 3.4. Food safety, sustainability, and cultural implications

Besides economic aspects, species mislabeling has implications for food safety, sustainability, and cultural practices. Some substitutions identified in this study involve species with dietary restrictions due to potential health risks. For instance, *L. flavobrunneum* (escolar), which we identified being sold as tilapia, contains high levels of indigestible wax esters that can cause severe gastrointestinal distress when consumed in large quantities (Fariñas Cabrero et al., 2015; Ho Ling et al., 2009; Pollack et al., 2018). Similarly, the mislabeling of *P. hypophthalmus* (striped catfish) as flounder is concerning, as pangasiids are known to accumulate heavy metals due to their farming conditions in the Mekong River, one of the world’s most polluted waterways (Ferrantelli et al., 2012; Nghia et al., 2009; Rodríguez et al., 2018). Consumers who believe they are purchasing one species may unknowingly ingest another with different allergenic properties, heavy metal levels, or foodborne risks, undermining informed decision-making and public health policies.

From a cultural perspective, species mislabeling can negatively impact religious dietary practices. For example, individuals following *kosher* dietary laws, such as those in the Jewish community, rely on accurate labeling to ensure that the food they consume agree with religious requirements (Blech, 2009; Lin et al., 2017). In this study, samples sold as flounder, a *kosher* species, were identified as *P. hypophthalmus* (striped catfish), which is not considered *kosher*. Such substitutions pose major concerns for consumers who depend on correct labeling for religious compliance, an often overlooked consequence of seafood fraud.

From a sustainability perspective, species mislabeling can exacerbate overfishing and compromise conservation efforts. While most species identified in this study are classified as Least Concern (LC) by the IUCN, wild populations of *P. hypophthalmus* are listed as Endangered (EN), despite extensive farming in Asia (IUCN, 2025). A similar pattern is observed in *Oreochromis spp*. (tilapia), where some species are classified as Critically Endangered (CR) (IUCN, 2025), although most tilapia in Brazil originate from aquaculture. Given these distinctions, commercial labeling should differentiate between farmed and wild-caught fish, and stricter trade regulations could prevent the sale of threatened species. Large-scale fishing operations often generate substantial bycatch—non-target species unintentionally caught alongside commercially valuable fish. To minimize financial losses, these species are frequently marketed under broad umbrella terms, masking their true identity and enabling the sale of overexploited or less desirable species (Almerón-Souza et al., 2018; French & Wainwright, 2022; Santa Brígida et al., 2024). Therefore, mislabeling facilitates illegal, unreported, and unregulated fishing, allowing endangered, low-stock, or bycatch species to enter in the market undetected (Kroetz et al., 2020; Pardo et al., 2018; Willette et al., 2017). Furthermore, it distorts consumer demand, potentially driving overexploitation of vulnerable species while concealing unsustainable fishing practices (Cooke et al., 2011; Kroetz et al., 2020).

The persistence of mislabeling highlights the need for stricter enforcement of seafood traceability regulations. Current labeling requirements in Brazil allow for broad commercial naming categories, making it easier for suppliers and restaurants to obscure species identity (Instrução Normativa MAPA N° 29, 2015). The lack of standardized nomenclature, both nationally and internationally, hampers the monitoring of seafood trade and enables fraudulent substitutions to persist. Implementing stricter controls, such as requiring scientific names alongside vernacular names and ensuring proper documentation of catch origin, would facilitate more effective regulation. Seafood fraud and unregulated fishing are global challenges that demand coordinated international responses. Initiatives similar to the U.S. Seafood Import Monitoring Program (SIMP), which tracks imports using scientific names and numeric taxonomic codes, could improve transparency and reduce the entry of mislabeled fish into markets (Steinkruger et al., 2025). Enhanced traceability “from boat to plate” would not only benefit environmental conservation but also ensure that consumers receive accurate information about the seafood they purchase. Given the crucial role of restaurants in the supply chain, they should be required to disclose the origin and identity of the fish they serve, empowering consumers to make informed choices based on health, economic, and sustainability considerations. Increasing consumer awareness is essential, as informed demand for correctly labeled seafood can drive industry-wide improvements and promote responsible consumption. Ultimately, consumers must actively question the origin of their seafood, demand transparency, and hold businesses accountable. Responsible consumption is not just a personal decision but a collective force capable of shaping industry practices and protecting the future of global fisheries.

## CRediT authorship contribution statement

**Fernanda Almerón-Souza:** Conceptualization, Formal Analysis, Investigation, Methodology, Validation, Visualization, Writing – original draft, Writing – review & editing; **Leonardo T. Gonçalves:** Formal Analysis, Methodology, Software, Visualization, Writing – original draft, Writing – review & editing; **Nelson J. R. Fagundes:** Conceptualization, Funding acquisition, Project administration, Resources, Supervision, Validation, Writing – review & editing.

## Acknowledgements

The authors would like to thank Lilian Viana Teixeira (UFMG), Eduardo Eizirik (PUCRS), and Filipe Michels Bianchi (UFRGS) for contributions in earlier versions of this manuscript. This work was supported by the Conselho Nacional de Desenvolvimento Científico e Tecnológico (CNPq) and Coordenação de Aperfeiçoamento de Pessoal de Nível Superior (CAPES). NJRF is supported by CNPq, grant number: 316900/2023–0.

## Data availability

Sequences are available in GenBank under accession numbers PV594474–PV594512 (fish DNA barcodes) and ####–#### (bacterial sequences).

## Supplementary Material

Table S1. Summary of BOLD Barcode ID results, displaying confidence values and the number of supporting database sequences for each identification.

Table S2. Output from RDP classifier, using the CO1 classifier as training set, including taxonomic assignments and corresponding bootstrap (BS) support values for each sequence.

Table S3. Purchase details of sushi samples used in the logistic regression analysis. Information includes sample number, restaurant code, sushi unit price (BRL), neighborhood of the sushi restaurant (in the city of Porto Alegre), average household income in the neighborhood (expressed in minimum wage units per month), and whether the sample was identified as mislabeled (true) or not (false).

